# Gut Microbiome Perturbation Affected Conjugative Transfer of Antimicrobial Resistance Genes

**DOI:** 10.1101/2024.08.03.605226

**Authors:** Gokhan Yilmaz, Maria Chan, Calvin Ho-Fung Lau, Sabrina Capitani, Mingsong Kang, Philippe Charron, Emily Hoover, Edward Topp, Jiewen Guan

**Affiliations:** Ottawa Laboratory-Fallowfield, Canadian Food Inspection Agency, Ottawa, Ontario, Canada; Ottawa Laboratory-Carling, Canadian Food Inspection Agency, Ottawa, Ontario, Canada; INRAE, University of Burgundy, University of Burgundy Franche-Comté, Dijon Cedex, France

**Keywords:** Antimicrobial resistance, plasmid transfer, gut microbiome, antibiotic treatments, *Salmonella*

## Abstract

The global spread of antimicrobial resistance genes (ARGs) poses a significant threat to public health. While antibiotics effectively treat bacterial infections, they can also induce gut dysbiosis, the severity of which varies depending on the specific antibiotic treatment used. However, it remains unclear how gut dysbiosis affects the mobility and dynamics of ARGs. To address this, mice were pre-treated with streptomycin, ampicillin, or sulfamethazine, and then orally inoculated with *Salmonella enterica* serovar Typhimurium and *S*. Heidelberg carrying a multi-drug resistance IncA/C plasmid. The streptomycin pre- treatment caused severe microbiome perturbation, promoting high-density colonization of *S*. Heidelberg and *S*. Typhimurium, and enabling IncA/C transfer from *S*. Heidelberg to *S*. Typhimurium and a commensal *Escherichia coli*. The ampicillin pre-treatment induced moderate microbiome perturbation, supporting only *S*. Heidelberg colonization and IncA/C transfer to commensal *E. coli*. The sulfamethazine pre-treatment led to mild microbiome perturbation, favoring neither *Salmonella spp*. colonization nor conjugative plasmid transfer. The degree of gut dysbiosis also influenced the enrichment or depletion of ARGs associated with mobile plasmids or core commensal bacteria, respectively. These findings underscore the significance of pre-existing gut dysbiosis induced by various antibiotic treatments on ARG dissemination and may inform prudent antibiotic use practices.

## Introduction

The global spread of antimicrobial resistance (AMR) genes (ARGs) among pathogenic bacteria constitutes a severe public health concern, posing a significant threat to the effective treatment of an expanding array of bacterial infections [1]. The excessive and inappropriate use of antimicrobials in animal production has led to the widespread prevalence of antimicrobial-resistant bacteria (ARB) in agricultural systems and the environment [2, 3, 4]. Numerous governments and organizations have adopted a One Health approach, striving to reduce the risk of AMR transmission to humans through food consumption and environmental exposure [5, 6, 7]. To realize this objective, it is crucial to understand the dynamics of ingested food- or water-borne ARB and the factors that might influence the potential dissemination of their ARGs to pathogenic and commensal bacteria within the host gut microbiota.

The gut microbiota is a community of microorganisms that coexist symbiotically within the host digestive tract, playing a vital role in maintaining host metabolic homeostasis, regulating the immune system, and influencing susceptibility to pathogens [8]. Additionally, it serves as a reservoir for diverse antimicrobial resistance genes and determinants, collectively referred to as the resistome [9]. Various factors, such as genetic background, diet, life style, and antibiotic treatment, can impact the composition of the gut microbiota and its resistome [9, 10]. Specifically, antibiotic treatment has the potential to disrupt the balance of the gut microbiome, reduce microbial diversity, foster the overgrowth of opportunistic pathogens, modify metabolic functions, and weaken the immune response [11].

Of greater concern, the use of antibiotics may facilitate the emergence of antibiotic-resistant bacteria in the gut. Predictions based on three-dimensional protein structures indicated that gut bacteria might harbor over 6000 uncharacterized AMR determinants, with the majority intrinsic to dominant commensals and seldom shared with bacterial pathogens [12]. However, recent metagenomics analysis demonstrated that mobile genetic elements played a crucial role in mediating ARG acquisition in gut commensals following broad-spectrum antibiotic treatment in a murine model [13]. Moreover, a systematic study evaluating the impact of 144 different antibiotics on gut bacteria revealed that β-lactam resistance among gut commensals was strain-specific and likely associated with horizontal gene transfer [14]. Given the variability in antibiotic treatments, encompassing specific antibiotics used, dosage, and duration of treatment, a more comprehensive understanding is essential to elucidate how antibiotics may influence the dissemination of ARGs within the gut microbiota.

Plasmid conjugation stands as a primary mechanism for horizontal gene transfer in bacteria, facilitating the exchange of genetic material through direct cell-to-cell contact between two bacterial cells. Conjugative plasmids play a significant role in the dissemination of ARGs in bacteria [15]. Through direct exposure, antibiotics may act as selective drivers for ARB, influencing the dynamics and efficiency of conjugation [16–18]. Moreover, through indirect pre-exposure, heavy antibiotic dosages (*e.g*. one oral dose of streptomycin at 1 g kg^-1^) can disrupt the gut microbiota, promoting the colonization and expansion of ARB and fostering conjugation [19, 20]. However, it remains unclear whether ARB are able to colonize and disseminate their ARGs within the gut microbiome that have been pre-exposed to clinical or veterinary uses of antibiotics. We hypothesized that the colonization of ARB and the subsequent ARG dissemination would be positively associated with the levels of microbiome perturbation induced by the pre-exposure to antibiotics. Thus, we utilized a murine model, administering a heavy dose of streptomycin, a clinical dosage of ampicillin, a veterinary dosage of sulfamethazine, or no antibiotic as a negative control. Subsequently, we infected the mice with *Salmonella enterica* serotype Heidelberg (*S*. Heidelberg) as a donor of a multi-drug resistance IncA/C plasmid, and *S.* Typhimurium as a recipient. The present study involved a comparative analysis of the dynamics of the donor, recipient, transconjugant, and selected ARGs and an integrase gene (*intI1*), alongside an examination of the gut microbial composition. The ARGs chosen herein included a sulfonamide resistance gene (*sul1*), a streptomycin resistance gene (*strA*), a beta-lactam resistance gene (*cfxA*), a macrolide, lincosamide, and streptogramin B resistance gene (*ermF*), and a tetracycline resistance gene (*tetQ*). The *sul1* and *strA* genes, carried by the IncA/C plasmid [18], served as representatives of introduced ARGs that were associated with mobile gene elements, whereas, the *cfxA*, *ermF* and *tetQ* genes were representatives of ARGs highly prevalent in gut commensal bacteria [21, 22]. The *intI1* gene, also carried by the IncA/C plasmid, was one of the key players mediating ARG dissemination [13].

## Materials and Methods

### Bacteria

*Salmonella enterica* serotype Heidelberg (SL-312) was isolated from a Canadian chicken farm. It contains a conjugative IncA/C plasmid, encoding the genes *aph(3)-Ia*, *aph(3)-Ib* or *strA*, *aph(6)-Id* or *strB*, *bla*_TEM-1B_, *bla_CMY-2_*, *dfrA1*, *floR*, *sul1*, *sul2*, and *tetA*, and *intI1*. This *S*. Heidelberg isolate exhibits resistance to amoxicillin/clavulanic acid, ampicillin, cefazolin, cefoxitin, cefpodoxime, ceftiofur, cephamycin, chloramphenicol, streptomycin, sulfamethizole, tetracycline, and trimethoprim-sulphamethoxazole [18]. In the present study, *S*. Heidelberg served as the donor of the multi-drug resistance IncA/C plasmid. The recipient, *Salmonella enterica* serotype Typhimurium (SL1344) contains a mobile IncQ plasmid, encoding the *strA*, *strB* and *sul2* genes. To facilitate recovery of *S*. Typhimurium, a spontaneous rifampicin-resistant mutant was generated. Briefly, *S.* Typhimurium was cultured overnight in Luria-Bertani (LB; Miller formulation, Difco, Fisher Scientific, Ottawa, ON, Canada) broth at 37 °C. A 1.0 ml overnight culture was pelleted, re-suspended in 100 µl LB broth, and spread on LB agar supplemented with 50 µg ml^-1^ rifampicin (LB-Rif). After 24 hours of incubation, resistant colonies were selected and sub-cultured on LB-Rif agar to generate and maintain a *S.* Typhimurium rifampicin-resistant mutant culture.

### *In vitro* conjugation

*In vitro* conjugation between *S.* Heidelberg (donor) and *S.* Typhimurium (recipient) was assessed in 10^-1^

× LB broth, following the method described by Laskey *et al*. [18]. The enumeration of the donor, recipient, and transconjugant bacteria was performed using XLT4 agar (Difco, Fisher Scientific, Ottawa, ON, Canada) supplemented with 50 µg ml^-1^ ampicillin (XLT4-Amp); 50 µg ml^-1^ rifampicin (XLT4-Rif); and 50 µg ml^-1^ ampicillin and 50 µg ml^-1^ rifampicin (XLT4-Amp-Rif), respectively. The frequency of conjugation is quantified as the ratio of transconjugant to donor bacteria counted at the end of the mating incubation period.

### *In vivo* conjugation

Mouse experiments and procedures, in compliance with guidelines and ethical standards of the Canadian Council on Animal Care and the ARRIVE guidelines, were approved by the Animal Care Committee at the Ottawa Laboratory-Fallowfield, Canadian Food Inspection Agency, Ottawa, Canada. Female C57BL/6 mice, 28-day old, were acquired from Charles River Laboratories (Saint Constant, QC, Canada). The mice were mixed and acclimatized for two weeks before receiving antibiotic treatments. They were then housed two or three per cage (Optimice^®^, Animal Care Systems, CO, USA) with water and feed provided *ad libitum*. A total of 21 mice were randomly divided into four groups for antibiotic pre-treatments: no antibiotic, ampicillin, streptomycin, and sulfamethazine (Table 1). Twenty-four hours after antibiotic withdrawal, the mice were first inoculated with the recipient and then with the donor bacteria one hour later. Bacterial inocula, consisting of 100 µl of log-phase culture with approximately 3.0 × 10^8^ colony forming units (CFU) of either the recipient or donor bacteria in phosphate buffered saline (PBS, pH 7.2), were administrated via oral gavage. The schedule of the experimental procedures is depicted in Figure 1. On designated sampling days, the mice were weighed and monitored for clinical symptoms like ruffled coat, hunched posture, and lethargy. Animals were euthanized upon development of morbidity, defined as showing clinical symptoms and/or experiencing a weight loss exceeding 20% from the day of bacterial inoculation. Fecal pellets were collected from all live mice on -7 (baseline), 0 (day of bacterial inoculation), 1, 3, 7, 10, 14, 17, and 21 days post-infection (DPI). These fecal pellets were processed as described by Laskey *et al.* [18] for DNA extraction and bacterial enumeration. The three selective agars, XLT4 agar supplemented with 50 µg ml^-1^ ampicillin and 50 µg ml^-1^ tetracycline (XLT4-Amp-Tet); 50 µg ml^-1^ rifampicin and 50 µg ml^-1^ streptomycin (XLT4-Rif-Strep); and 50 µg ml^-1^ ampicillin, 50 µg ml^-1^ rifampicin, and 50 µg ml^-1^ streptomycin (XLT4-Amp-Rif-Strep), were used to enumerate the donor, recipient and putative *S*. Typhimurium transconjugant, respectively. The limit for bacterial enumeration was 2.2 log_10_ CFU g^-1^ in feces. Additionally, Chromocult agar (EMD Millipore, Toronto, ON, Canada) supplemented with 50 µg ml^-1^ ampicillin (Chr-Amp) was used to isolate potential commensal *Enterobacteriaceae* transconjugants.

**Figure 1.**
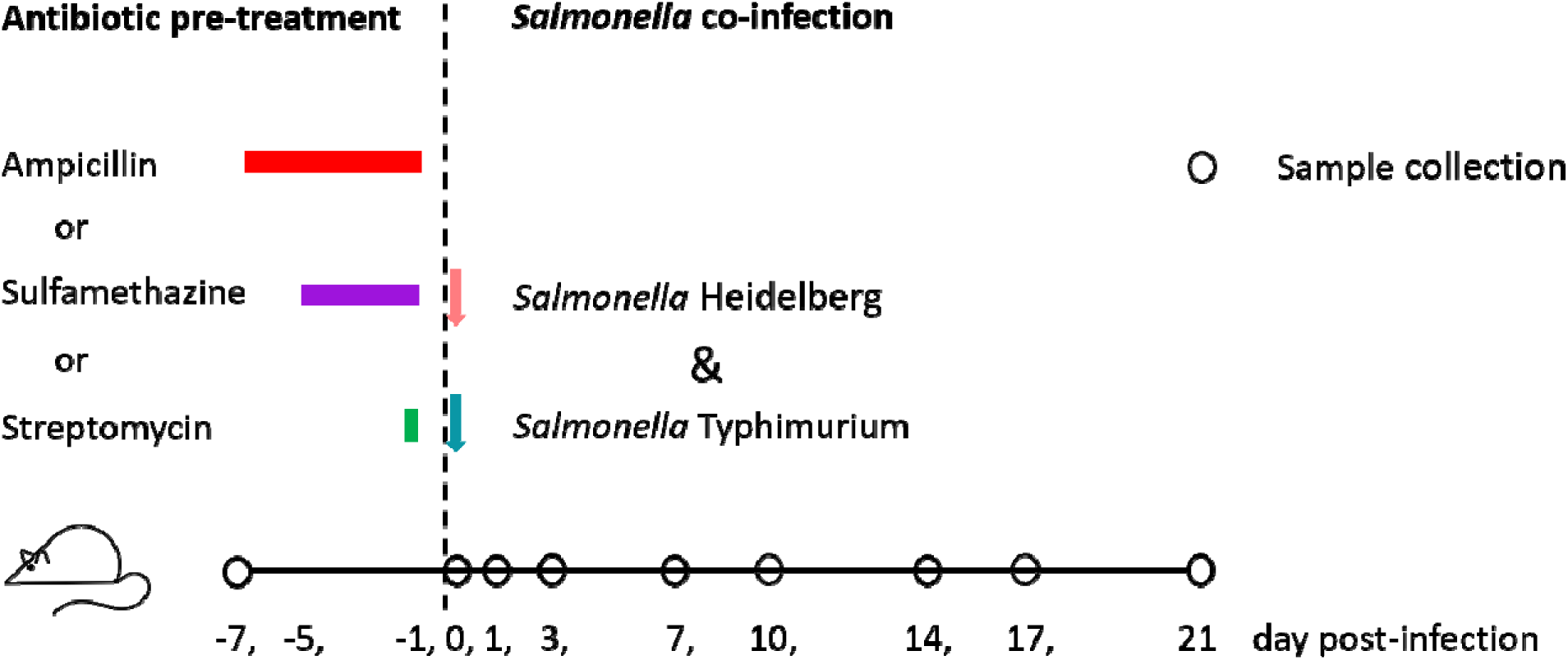
Schedule of procedures for mice inoculated with *Salmonella* Typhimurium (recipient) and *Salmonella* Heidelberg (donor) following pre-treatment with either ampicillin, sulfamethazine, streptomycin, or no antibiotic. Sample collection was performed on various days post-infection (DPI) as indicated. Colored bars represent the duration of antibiotic pre-treatments, and arrows indicate the points of bacterial inoculation during the co-infection phase.

**Table 1.** Treatment groups in mouse experiments.

| Group | Antibiotic pre-treatment<sup>*</sup> | n |
| --- | --- | --- |
| None | Control (no antibiotic) | 5 |
| Amp | Ampicillin, 40 mg kg<sup>-1</sup> d<sup>-1</sup> via drinking water for 7 days | 6 |
| Strep | Streptomycin, one dose of 20 mg per mouse via oral gavage | 5 |
| Sulf | Sulfamethazine, 225 mg kg<sup>-1</sup> d<sup>-1</sup> via drinking water for 5 days | 5 |

<sup>\*</sup> Antibiotic pre-treatment was stopped 24 hours before bacterial inoculation in all cases.

### Whole genome sequencing

To identify putative transconjugant bacteria, colonies displaying distinct morphologic characteristics on XLT4-Amp-Rif-Strep and Chr-Amp media (3 colonies each morphologic type) were isolated for each mouse. These isolates were sub-cultured on LB agar to generate single-isolated colonies. DNA extraction from these isolates followed the method described by Laskey *et al*. [18]. The extracted DNA was subjected to polymerase chain reactions (PCRs) using primers for *invA*, *bla_CMY-2_*, and *bla_TEM-1_* genes (Table S1). Isolates that tested negative for the *invA* gene but positive for both *bla_CMY-2_*, and *bla_TEM-1_* genes were selected for whole-genome sequencing (WGS). Isolates of the *S.* Heidelberg donor and *S*. Typhimiurium recipient and some ampicillin-resistance bacteria were sequenced as well. The WGS was performed on the Illumina MiSeq system (Illumina Canada, Vancouver, BC, Canada). The library preparation was performed using the Illumina DNA Prep Kit (Illumina). This process yielded paired-end reads of 300 base pairs (bp). Genome assembly and analysis was performed using COWBAT v0.5.0.23 (https://github.com/OLC-Bioinformatics/COWBAT), a comprehensive tool integrating several steps such quality control (QC) trimming with BBMap v38.96 (https://sourceforge.net/projects/bbmap/), assembly with SKESA v2.1 [23], quality assessment with QUAST v5.1.0 [24], and plasmid identification with MOB-suite v 3.0.3 [25].

### 16S rRNA gene amplicon sequencing

DNA samples extracted from mouse fecal pellets using the NucleoSpin Soil DNA Extraction Kit (Macherey-Nagel, Germany), underwent 16S rRNA gene amplicon sequencing. Briefly, the V3-V4 region of the 16S ribosomal RNA gene was amplified using PCR [26]. The sequencing libraries were prepared using the Nextera XT Index Kit v2 (Illumina), and then purified and normalized with the NGS Normalization 96-well kit (Norgen Biotek, Thorold, ON, Canada).

Sequencing was performed on the Illumina MiSeq system using the MiSeq v3 kit with a 10% PhiX spike-in, targeting an output of 100,000 raw reads per sample. QC filtration of the raw reads was performed using Fastp v 0.23.2 [27], and primer sequences were removed using pTrimmer v 1.3.4 [28]. The processed reads were then classified with Emu v 3.4.5 with the pre-built Emu database for accurate microbial identification [29]. Data analysis and visualization were performed using the following R packages: vegan v 2.6-2, ggplot2 v 3.3.6, phyloseq v 1.38.0, and microbiomeMarker v 1.02.

### Quantification of antimicrobial resistance and integrase genes

The abundance of *strA*, *sul1*, *intI1*, *cfxA*, *emrF*, *tetQ*, and the small subunit ribosomal RNA fragment 1 gene (*rrnS1*) was determined using quantitative polymerase chain reaction (qPCR) with a QuantStudio 3 Real-Time PCR System (Thermo Fisher, Nepean, ON, Canada). The reaction mixture, with a total volume of 12.5 µl, contained 6.25 µl of Power SYBR Green PCR master mix (Thermo Fisher), 0.625 µl each of forward and reverse primers (primer sequences and final concentrations detailed in Table S1, 2.5 µl of template DNA normalized to 0.4 ng µl^-1^, and 2.5 µl of nuclease free water. The PCR program included an initial incubation at 95°C for 10 minutes, followed by 40 cycles of denaturation at 95°C for 15 seconds, with annealing and elongation set at specific temperatures and times (Table S1). A final melting curve analysis, ramping the temperature from the elongation setting to 95°C, verified the specificity of the PCRs. Each DNA sample underwent triplicate PCR reactions. Standards for qPCR were prepared by synthesizing the sequences of all target amplicons and cloning them into a pUC-IDT plasmid (IDT, Coralville, Iowa, USA). The recombinant plasmid was transformed into an *Escherichia coli* DH-5α strain for amplification, and then extracted from the bacterial cells. The plasmid DNA was linearized, quantified, and used as standards for the construction of the qPCR standard curves. Detection limits for the various gene targets were 4.3 ± 0.6 log_10_ copies per gram of feces.

### Statistical analyses

Differences in conjugation frequency, mean abundance of each target bacterium, fold change of each targeted gene, and the relative abundance of each phylum, family, or genus in the 16S rRNA gene community profiles between treatment and control groups on identical sampling days were analysed with Welch’s ANOVA followed by Dunnett’s test, using GraphPad Prism 8.0 software. All correlations were tested using the Pearson correlation test. The treatment groups contained 5 or 6 mice (Table 1), and the mean value derived from technical replicates of a fecal pellet from each mouse on every sampling day constituted one data point. A *p*-value < 0.05 was considered statistically significant. Linear discriminant analysis (LDA) effect size (LEfSe) [30] was used to identify the taxa that best discriminated one treatment group from the others on 0 day post-infection.

## Results

### *Salmonella* colonization and plasmid conjugation

In mice co-infected with *S*. Heidelberg (donor) and *S*. Typhimurium (recipient), fecal shedding of both bacterial strains showed similar patterns at most post-infection time points within each antibiotic pre- treatment group (Fig. 2 A-D). Both strains peaked at 1 DPI and gradually decreased to undetectable levels in most mice over the 21 days post-infection. Notably, mice pre-treated with streptomycin exhibited a significantly higher abundance of both *Salmonella* strains compared to the control that received no antibiotic pre-treatment (Fig. 2 A-D, Fig. S1). On 1 DPI, both *S*. Heidelberg and *S*. Typhimurium reached a peak abundance of 7.4 log_10_ g^-1^, while the *S*. Typhimurium transconjugant reached 3.9 log_10_ g^-1^. The observed *in vivo* conjugation frequency, 6.5 (± 3.3) ×10^-4^, was much higher than the *in vitro* conjugation frequency, 4.9 (± 3.7) ×10^-6^ (*p* < 0.05). Additionally, *Escherichia coli* (*E. coli*) transconjugants were recovered from these mice (Table 2). In mice pre-treated with ampicillin, *S*. Heidelberg showed a peak abundance at 5.6 log_10_ g^-1^, while *S*. Typhimurium did not exceed 3.8 log_10_ g^-1^. In this group, only *E. coli* transconjugants were detected, with no *S*. Typhimurium transconjugants recovered (Table 2). In contrast, mice that received sulfamethazine or no antibiotic pre-treatment maintained *Salmonella* strains at levels below 4.3 log_10_ g^-1^ throughout the 21-day post-infection period, with no transconjugants being recovered. Isolates of the donor, recipient, transconjugants, and some ampicillin-resistance bacteria underwent WGS. The sequencing data confirmed that the IncA/C plasmid was transferred from the *S*. Heidelberg donor to the *S*. Typhimurium recipient and the commensal *E. coli*. Additionally, the sequencing data confirmed the presence of the IncQ plasmid, encoding the *strA*, *strB* and *sul2* genes, in *S*. Typhimurium (SAMN40034283), and also showed intrinsic ARGs in some ampicillin-resistance commensal bacteria, such as the *mdfA* and *bla_ACT_*genes in the chromosomes of *E. coli* (SAMN40034292) and *Enterobacter xiangfangensis* (SAMN40034293), respectively.

**Figure 2.**
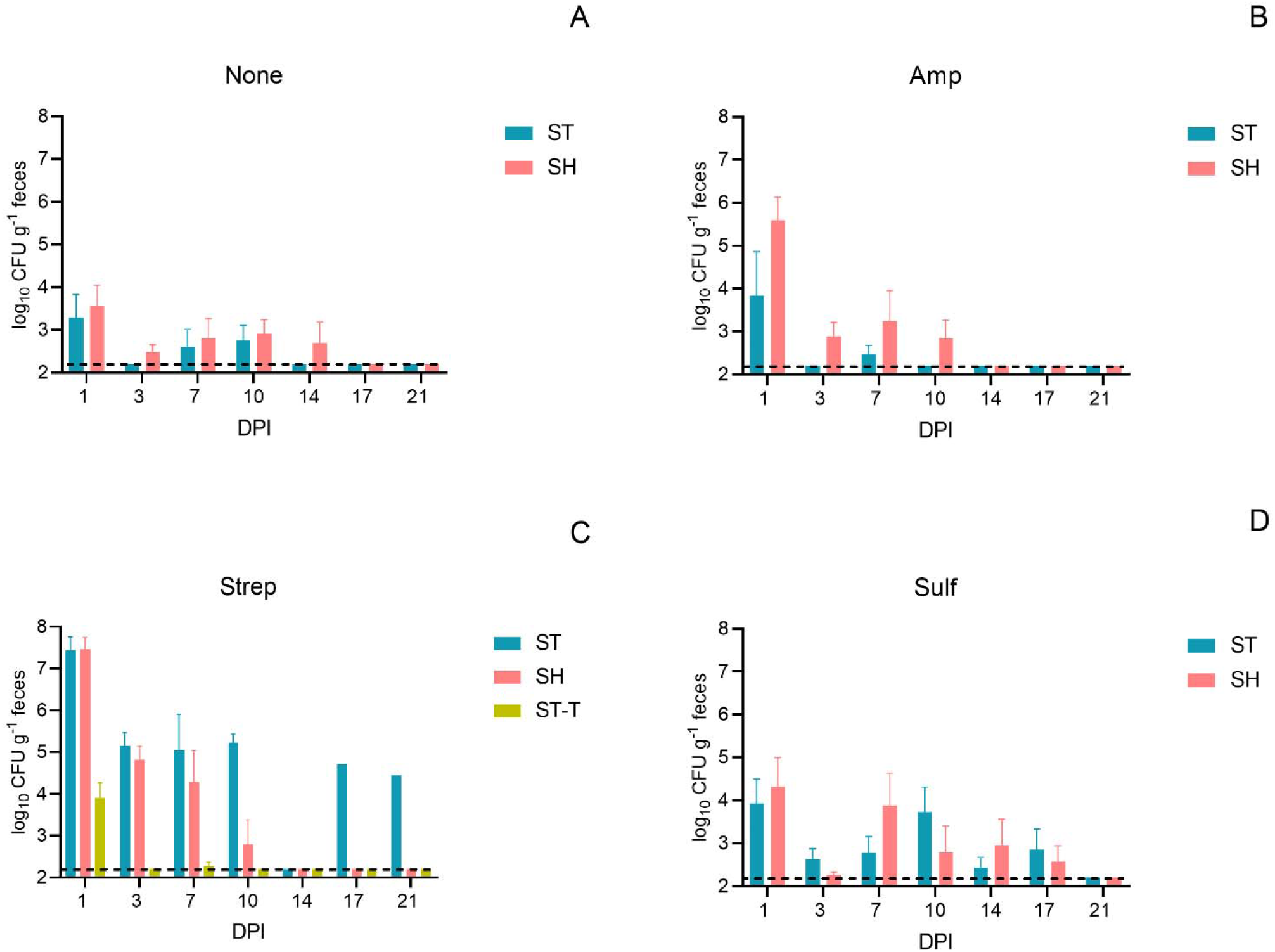
Abundance (mean ± SE) of *Salmonella* Heidelberg (SH, donor), *S*. Typhimurium (ST, recipient) and *S.* Typhimurium transconjugants (ST-T) in fecal samples from mice that received both the recipient and donor inoculation following different pre-treatments. The pre-treatments include no antibiotic (None, A), ampicillin (Amp, B), streptomycin (Strep, C), and sulfamethazine (Sulf, D). The y-axis shows the logarithmic scale of colony-forming units per gram of feces (log_10_ CFU g^-1^), and the x-axis denotes the days post-infection (DPI). The dash lines represent the detection limit of 2.2 log_10_ CFU g^-1^.

**Table 2.** Transconjugants isolated from mice subjected to antibiotic pre-treatments followed by Salmonella infection.

| Pre-treatment | Transconjugant isolated<br>(day post-infection)* | Accession number** |
| --- | --- | --- |
| None | Not detected |  |
| Amp | *Escherichia coli* (1) | SAMN40034289 - SAMN40034292 |
| Strep | *Salmonella* Typhimurium (1, 7) | SAMN40034283 - SAMN40034288 |
|  | *Escherichia coli* (1, 3, 7) | SAMN40034294 - SAMN40034303 |
| Sulf | Not detected |  |

\* All transconjugants containing the IncA/C plasmid were confirmed via whole genome sequencing.
\*\* BioSample accession number of representative isolates

### Mouse survival post *Salmonella* infection

All mice that received ampicillin or no antibiotic pre-treatment survived for a minimum of 21 days following co-infection with *S.* Typhimurium and *S.* Heidelberg (Fig. S2). In the group pre-treated with sulfamethazine, one out of five mice was dead on 3 DPI, while the remaining four mice survived the 21-day post infection period. In contrast, survival rates were significantly lower (*p* < 0.05) among mice that received the streptomycin pre-treatment when compared to the other groups. Specifically, four out five mice with streptomycin pre-treatment were found dead or euthanized due to morbidity, one on 3 DPI, one on 10 DPI and two on 14 DPI. Only one mouse in the streptomycin pre-treatment group survived the 21-day post infection period.

### Dynamics of antimicrobial resistance and integrase genes

The antibiotic pre-treatments did not cause direct impacts on the dynamics of *strA*, *sul1*, and *intI1*genes, that were carried by the IncA/C plasmid. On 0 DPI, the abundance of these three genes remained unchanged across all mice, regardless of the pre-treatment (Fig. 3 A-C). However, following co-infection with the *Salmonella* donor and recipient, a noticeable increase in the abundance of all three genes was observed on 1 DPI in most mice. Mice that received streptomycin pre-treatment showed a significant enrichment of these genes (*p* < 0.05) compared to those without antibiotic pre-treatment from 1 to 7 DPI. Furthermore, the abundance of the *sul1* and *intI1* genes was positively correlated with that of *S*. Heidelberg (Pearson’s correlation coefficient *r_sul1_* = 0.7229, *r_intI1_* = 0.6580, Fig. S3). The elevated levels of *sul1* and *intI1* genes observed initially declined by 21 DPI across all mice (Fig. 3 B & C). The abundance of the *strA* gene was positively correlated with that of *Salmonella* spp. (*r_strA_* = 0.5816, Fig. S3), as both donor and recipient carry the *strA* gene. The *strA* abundance remained elevated in streptomycin pre-treatment group due to the persistence of *S.* Typhimurium for the 21-day post infection period (Fig. 3 A and 2 C).

**Figure 3.**
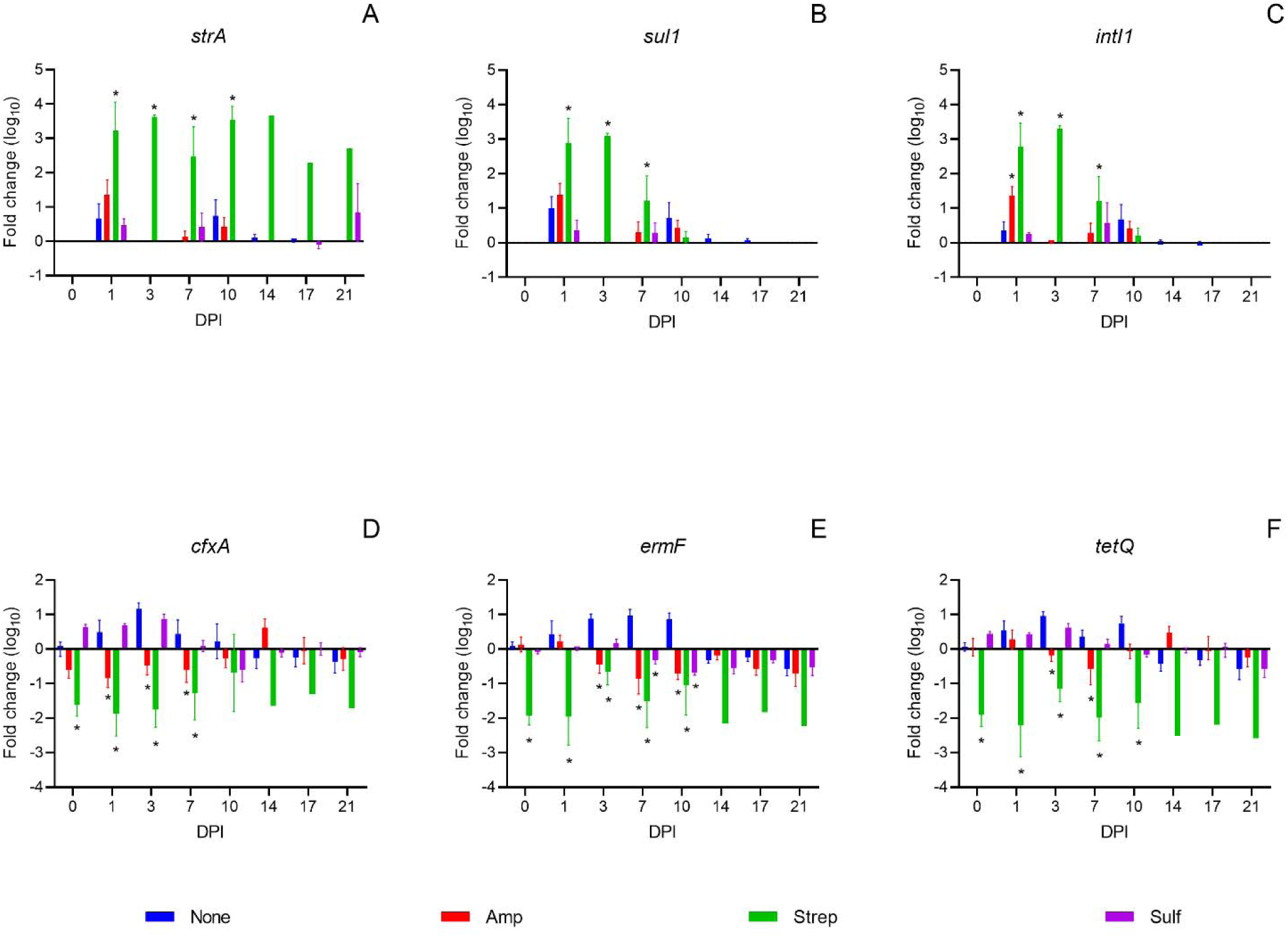
Fold change in the abundance of antimicrobial resistance genes *strA*, *sul1*, *cfxA*, *tetQ,* and *ermF*, and an integrase gene, *intI1*, (mean + SE) in fecal samples from mice following inoculation with *Salmonella* Typhimurium and *Salmonella* Heidelberg. Mice were pre-treated with either no antibiotic (blue), ampicillin (red), streptomycin (green) or sulfamethazine (purple). Gene abundances were averaged from triplicate technical replicates and were normalized against that of the *rrnS1* genes for each mouse. Fold changes from 0 to 21 days post-infection (DPI) were calculated relative to the baseline on -7 DPI. Asterisks (*) indicate a significant difference (*p* < 0.05) between an antibiotic pre-treatment group and the control group with no antibiotic pre-treatment within each DPI, as determined by Welch’s ANOVA followed by Dunnett’s test.

In comparison, the antibiotic pre-treatments caused direct impacts on the dynamics of *cfxA*, *ermF* and *tetQ* genes, the representative ARGs carried by commensal bacteria. The abundance of these three genes significantly (*p* < 0.05) decreased on 0 DPI, and the decrease remained to 10 DPI in mice receiving the streptomycin pre-treatment (Fig. 3 D-F). In mice pre-treated with ampicillin and sulfamethazine, the *cfxA* gene showed minor fluctuations, while the *ermF* and *tetQ* genes were relatively stable on 0 DPI. However, post *Salmonella* co-infection, the abundance of *cfxA*, *ermF* and *tetQ* genes significantly decreased on various DPIs in mice receiving the ampicillin pre-treatment. Whereas, in the sulfamethazine treatment group, the abundance of these genes fluctuated in a way similar to those in mice receiving no antibiotic pre-treatment (Fig. 3 D-F).

### Association between the gut microbiome and the dynamics of target genes

To investigate the association between the gut microbiome and the dynamics of target genes, the taxonomic composition of gut microbial communities was analyzed using 16S rRNA gene amplicon sequencing. All three antibiotic pre-treatments significantly reduced the species richness within the gut microbiome compared to the control, with the streptomycin pre-treatment caused the greatest reduction, followed by the ampicillin and the sulfamethazine pre-treatments (Fig. 4 A). The dissimilarity of the gut microbiome between each treatment group and the control also reflected the different levels of perturbation induced by the antibiotic pre-treatments (Fig. 4 B). The gut microbiome gradually recovered by 7 DPI in mice receiving the ampicillin and the sulfamethazine pre-treatment, but not the streptomycin pre-treatment (Fig. S4, S5). The antibiotic pre-treatments caused differential impacts on the microbial composition. The relative abundance of Proteobacteria increased on 0 DPI following the ampicillin and streptomycin pre-treatments compared to the no antibiotic control (Fig. 4 C). Post *Salmonella* infection, the relative abundance of Proteobacteria continued to rise with this increase persisting until at least 7 DPI in mice that received the streptomycin pre-treatment. In comparison, the relative abundance of Proteobacteria in mice that received the ampicillin pre-treatment returned to levels comparable to the no antibiotic control group on 7 DPI. The relative abundance of Proteobacteria in mice pre-treated with sulfamethazine was similar to that of the no antibiotic control group from -7 to 7 DPI. Furthermore, the relative abundance of *Escherichia* and Enterobacteriaceae increased following streptomycin and ampicillin pre-treatment on 0 and 1 DPI (Fig. 4 E, F).

**Figure 4.**
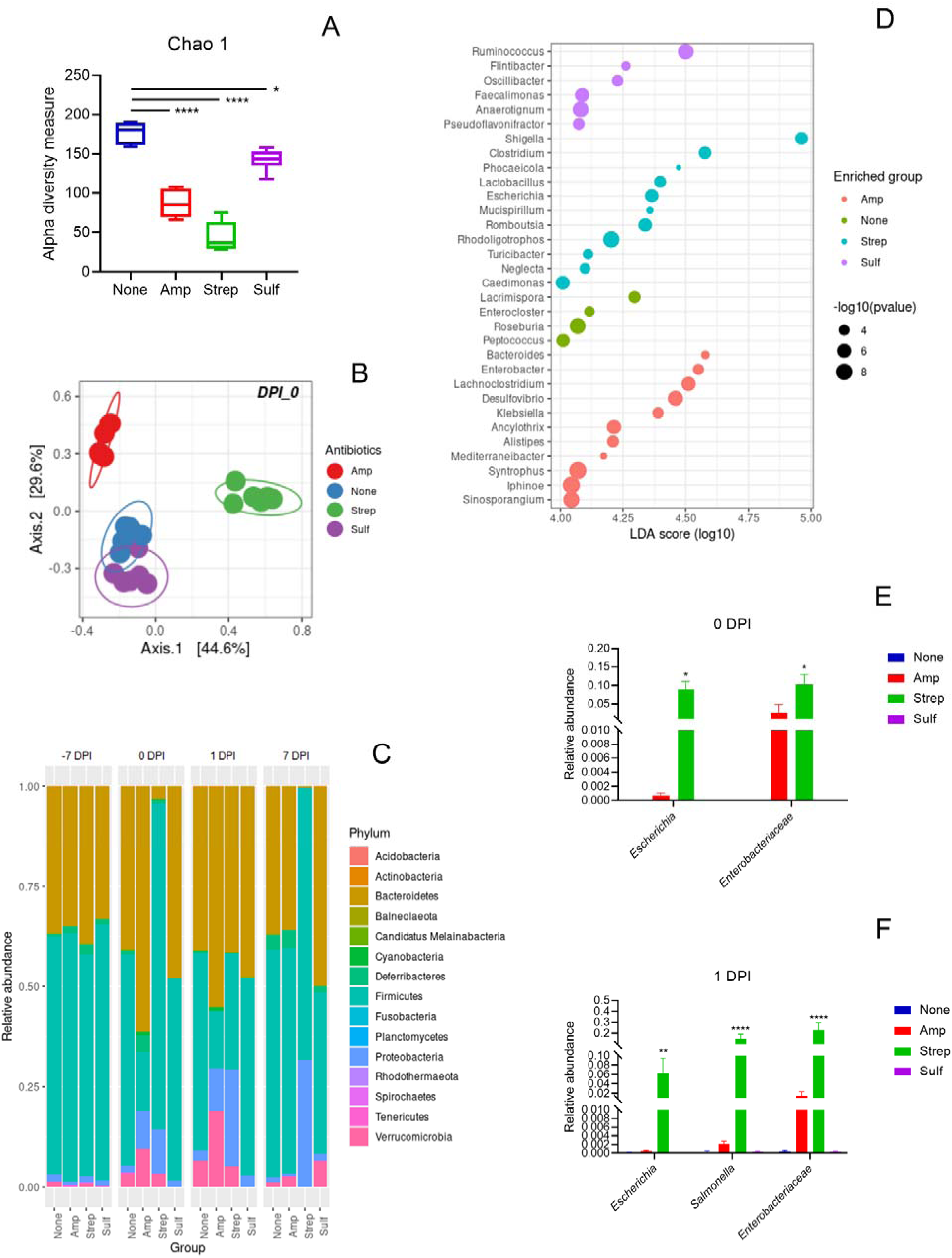
Analysis of the mouse gut microbiome using 16S rRNA gene amplicon sequencing: (A) microbial species richness assessed by the Chao1 index on 0 day post-infection (DPI) following various antibiotic per-treatments: None (no antibiotic, blue), Amp (ampicillin, red), Strep (streptomycin, green), Sulf (sulfamethazine, purple), with statistical significance denoted by asterisks (\**p* < 0.05, \*\*\*\**p* < 0.0001). (B) Principal coordinate analysis (PCoA) based on the Bray-Curtis dissimilarity of the gut microbiome on 0 DPI, with colors indicating different antibiotic treatments. (C) Bar graph showing the mean relative abundance of microbial phyla from -7 to 7 DPI across different treatment groups. (D) Dot plot illustrating the differential enrichment of various genera on 0 DPI, with dot size representing the negative logarithm of the *p*-value and color indicating the antibiotic treatment group. (E-F) Relative abundance of Enterobacteriaceae on 0 (E) and 1 (F) DPI, with bars representing the mean values and error bars indicating standard error (SE). Statistical significance in panels E and F is indicated by asterisks above the bars (\*\**p* < 0.01, \*\*\*\**p* < 0.0001).

The differential-abundance analysis showed that various genera were enriched following the different antibiotic pre-treatment. After ampicillin pre-treatment, genera such as *Alistipes*, *Bacteroides*, *Enterobacter*, *Lachnociostridium* and *Klebsiella* showed enrichment. Control mice with no antibiotic pre-treatment exhibited an increase in *Enterocloster*, *Lacrimispora*, *Peptococcus* and *Roseburia.* The streptomycin pre-treatment resulted in an enrichment of *Clostridium*, *Escherichia*, *Phocaeicola*, *Rhodoligotrophos*, *Romboutsia* and *Shigella*. Sulfamethazine pre-treatment led to an increase in *Anaerotignum*, *Faecalimonas*, *Ruminoccoccus* and *Pseudoflavonifractor* (Fig. 4 D). Among these enriched genera, *Alistipes*, *Bacteroides* and *Phocaeicola* were found to be positively associated with the the *cfxA*, *ermF* and *tetQ* genes, while *Escherichia*, *Romboutsia* and *Shigella* were negatively associated with these genes (Fig. 5). In addition, the Bacteroidetes phylum was positively, and the Proteobacteria phylum negatively associated, with the *cfxA*, *ermF*, and *tetQ* genes. The Proteobacteria phylum, along with the *Enterobacteriaceae* family and the *Escherichia* and *Shigella* genera, were positively associated with the *strA*, *sul1*, and *intI1*genes (Fig. 5). Within the Firmicutes phylum, the Lachnospiraceae family was associated positively with the *cfxA*, *ermF* and *tetQ* genes and negatively with the *strA* gene, whereas the Peptostreptococcaceae family showed the opposite trend.

**Figure 5.**
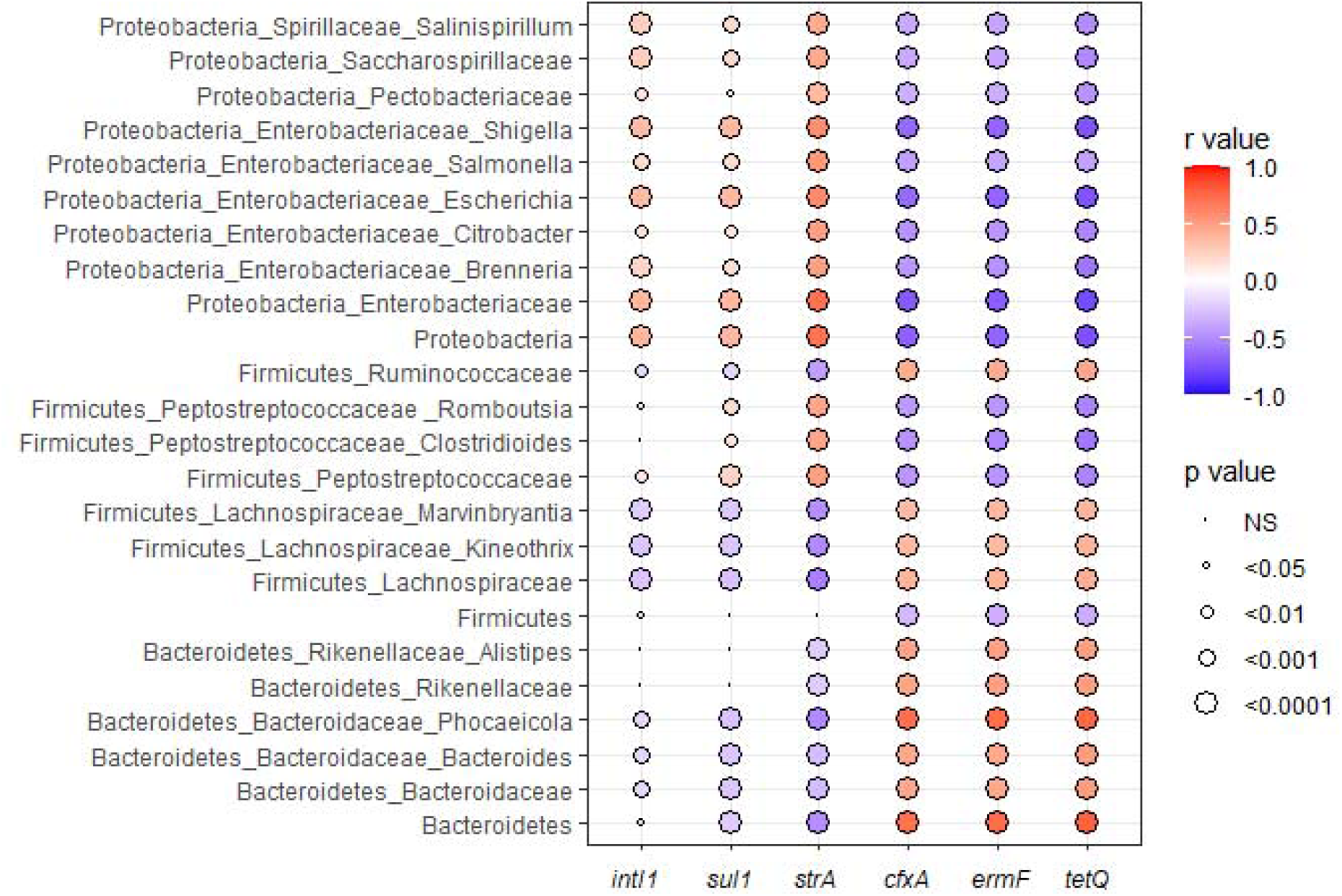
Correlogram shows Pearson’s correlations (*r* values) between the abundance of an integrase gene (*intI1*) and specific antimicrobial resistance genes (*sul1*, *strA*, *cfxA*, *ermF*, and *tetQ*), and various bacterial taxa at different taxonomic levels (phylum, family and genus) in the fecal samples of all mice in this study. The color intensity of each dot corresponds to the strength and direction of the correlation, with read indication a positive correlation and blue a negative correlation. The size of the circles represents the level of statistical significance, with larger circles indicating a lower *p*-value. Non-significant correlations are marked as ‘NS’.

## Discussion

Antibiotics are powerful medications that treat bacterial infection, but they also drive the dissemination of ARGs within the gut microbiome. The presence of antibiotics grants a selective advantage for ARB to colonize and replicate in the gut, enabling the conjugative transfer of resistance plasmids from the ARB to opportunistic pathogens and commensals, and subsequently encouraging the transconjugant propagation [17, 18]. Furthermore, antibiotics collaterally target commensals, altering the gut microbiome composition and causing dysbiosis [31]. As such, immediately following antibiotic treatment, the gut microbiome may provide a great ecological opportunity for ARB to colonize and disseminate their ARGs, even in the absence of antibiotics [32, 33]. Previous studies have demonstrated the conjugative transfer of antibiotic resistance plasmids following heavy-dose streptomycin treatments for clearing the mouse gut [19, 20]. Depending on the chemical composition, formulation and administration route, antibiotics may cause different magnitudes of perturbation to the gut microbiome [34]. Despite reports on plasmid transfer related to clinical antibiotic treatment [35, 36], a deeper understanding of how microbiome disturbances facilitate the conjugative transfer of antibiotic resistance plasmids is needed.

In this study, pre-treatments with a heavy-dose of streptomycin, a clinical dosage of ampicillin and a veterinary dosage of sulfamethazine were used to generate different levels of microbiome perturbation, along with no antibiotic pre-treatment as a negative control. All three antibiotic pre-treatments significantly reduced the species richness in the gut microbiome compared to the control. The streptomycin pre-treatment caused a severe perturbation with the greatest reduction in species richness, depleting the Bacteroidetes and enriching the Proteobacteria phyla. The disturbed gut microbiome facilitated the high-density colonization of both the *Salmonella* donor and recipient (> 7 log_10_ CFU g^-1^), and promoted the transfer of the multi-drug resistance IncA/C plasmid from the *Salmonella* donor to the *Salmonella* recipient via conjugation. The intensive *Salmonella* infection and severe gut dysbiosis might contribute to the significantly lower survival rate observed in mice pre-treated with streptomycin. In contrast, the clinical dosage of ampicillin caused a moderate microbiome perturbation with a medium reduction in species richness, depleting the Firmicutes and enriching the Proteobacteria phyla, which supported the colonization of the *S.* Heidelberg donor (5 to 6 log_10_ CFU g^-1^) but not the *S.* Typhimurium recipient. Thus, the conjugation between the *Salmonella* donor and recipient was not detected following the ampicillin pre-treatment. However, both streptomycin and ampicillin pre-treatments enriched the Enterobacteriaceae family. Within this family the *Escherichia* and *Shigella* genera were significantly enriched by the streptomycin pre-treatment, and the *Enterobacter* and *Klebsiella* genera by the ampicillin pre-treatment. Although the relative abundance of *Escherichia* was lower following the pre- treatment with ampicillin than streptomycin, *E. coli* served as a commensal recipient and supported the dissemination of the IncA/C plasmid in the gut microbiome that was disturbed by either pre-treatment. In support of our findings, Stecher *et al*. [37] reported that pathogen-driven inflammatory responses generated a transient expansion of Enterobacteriaceae which promoted horizontal gene transfer via conjugation between pathogens and commensals in the mouse gut. In comparison, the sulfamethazine pre-treatment only caused a mild microbiome perturbation with the least reduction in species richness, which did not favor the colonization of the *Salmonella* donor or recipient, or the conjugative transfer of the IncA/C plasmid. This finding aligns with the report by Liu *et al*. [38] on the negligible impact of sulfamethoxazole on the abundances of Bacteroidetes and Firmicutes in the mouse gut. In our control mice receiving no antibiotic pre-treatment, the normal gut microbiome resisted *Salmonella* colonization and eliminated IncA/C transfer. Overall, our data suggest that the colonization of opportunistic antimicrobial resistance food- or water-borne pathogens and the subsequent dissemination of the introduced ARGs are positively associated with the perturbation magnitude of the gut microbiome and the expansion of Enterobacteriaceae.

Furthermore, various antibiotics may induce differential changes to the structure of the gut microbiome and drive the dissemination of ARGs in the enriched taxa. Using a C57BL/6J mouse model, de Nies *et al*. [13] reported that the relative abundance of ARGs, including those from the beta-lactam, glycopeptide, and aminoglycoside categories, was significantly increased in the enriched Akkermansiaceae family in the gut microbiome after treatment with an antibiotic cocktail of ampicillin, vancomycin, neomycin, and metronidazole. They suggested that integrons associated with ARGs played a key role in mediating the AMR spread. Similarly, Xu *et al*. [40] observed that mono-antibiotic treatment with ampicillin, ciprofloxacin, or fosfomycin, led to an increased relative abundance of specific bacterial species and ARGs in Balb/c mice. They suggested that the enrichment of transposases after treatment with ciprofloxacin could signify an increased potential for horizontal gene transfer in the gut microbiome. Furthermore, a study involving human participants by Anthony *et al.* [31] reported significantly increased ARG burden after treatments with cefpodoxime, azithromycin, or a combination of both, indicating antibiotic-specific changes in ARG relative abundance. Despite these insights, a common finding across all these studies [13, 31, 40] was a significant reduction in species richness and the relative abundance of numerous ARGs after antibiotic treatments. In accordance with these studies, we observed that the species richness of the gut microbiome was significantly reduced following each of the three antibiotic pre-treatments, accompanied by the significant enrichment of specific genera depending on the antibiotic used. We found a significant enrichment of spore-forming bacteria and members of the Bacteroidaceae, Rikenellaceae and Enterobacteriaceae families. Likely, these bacteria can tolerate antibiotics through spore-mediated persistence, intrinsic resistance and/or other resiliency mechanisms [34]. In our study, the commensal *E. coli* carried a *mdfA* gene in its chromosome. The *mdfA* is a multi-drug transporter gene conferring resistance to certain antibiotics such as chloramphenicol, erythromycin, and certain aminoglycosides and fluoroquinolones [40]. Similarly, we isolated a commensal *Enterobacter* strain carrying a *bla_ACT_* gene in its chromosome. These intrinsic resistance determinants likely contributed to the enrichment of *Escherichia* and *Enterobacter* following the streptomycin and ampicillin pre-treatments, respectively. Positively associated with the Enterobacteriaceae enrichment, the abundance of the *strA* and *sul1* genes increased after the colonization of the *Salmonella* donor carrying the IncA/C plasmid encoding these two genes. Clearly, the dissemination of *strA* and *sul1* genes in enriched Enterbobacteriaceae via the conjugative transfer of the plasmid was affected by the magnitude of dysbiosis that was induced by the different antibiotic pre- treatments. The dynamic of the *intl1* gene was similar to that of the *strA* and *sul1* genes, influenced similarly by the magnitude of gut dysbiosis. In support of our findings, Xu *et al*. [40] observed only a slight, short increase in the relative abundance of integrases in the mouse gut microbiome after ampicillin treatment. In contrast to the introduced ARGs, the abundance of the *cfxA*, *tetQ*, and *ermF* genes was reduced in the disturbed gut microbiome. The reduction varied in extent corresponding to the level of dysbiosis induced by the antibiotics, with the greatest decrease following streptomycin, a lesser decrease after ampicillin, and the smallest to no decrease post-sulfamethazine treatment. As the *cfxA*, *tetQ*, and *ermF* genes are highly prevalent in the Bacteroidetes and Firmicutes phyla [21, 22], the depletion of these core commensal bacteria by antibiotic treatment alongside *Salmonella* infection likely led to the reduction of these ARGs. Overall, our findings suggest that the dynamics of ARGs are closely connected to and largely affected by the taxonomic changes in the gut microbiome.

## Limitation

The use of conventional selective bacterial culture methods in this study provided effective isolation and enumeration of *S*. Typhimurium transconjugants. However, these techniques had limitations when attempting to identify and recover unknown transconjugants among the commensal bacteria. Due to resource constraint, our study only focused on recovering Enterobacteriaceae transconjugants and was not able to capture any potential ARG disseminations in other taxa. To investigate the dynamics of ARGs, we used qPCRs assays targeting specific ARGs known to be associated with mobile genetic elements or inherently present in core commensal bacteria. Such an approach clearly captured the interactions between the dynamics of target ARGs and the perturbations of the gut microbiome, but was not able to cover the broader resistome changes.

## Conclusion

This study explored the impacts of gut dysbiosis induced by various antibiotic pre-treatments on the conjugative transfer and the dynamics of ARGs using a mouse model. We found a positive correlation between the potential for conjugative transfer of a multi-drug resistance plasmid, from the *S.* Heidelberg donor to the *S*. Typhimurium recipient and the commensal *E. coli*, and the degree of gut dysbiosis, even in the absence of antibiotics providing a direct selection advantage. Furthermore, the magnitude of gut dysbiosis also affected the dynamics of ARGs. The increase in the abundance of the *sul1* and *strA* genes carried by the multi-drug resistant plasmid was positively associated with the enrichment of Proteobacteria and the Enterobacteriaceae, whereas, the reduction in the abundance of the *cfxA*, *tetQ*, and *ermF* genes was positively associated with the depletion of Bacteroidetes and Firmicutes. Our findings underline the importance of pre-existing gut dysbiosis induced by specific antibiotics on the horizontal transfer of ARGs from food- or water-borne ARB to commensals, and may help guiding antibiotic treatment choices to minimize the dissemination of AMR in the gut microbiome.

## Acknowledgements

Great appreciation is expressed to Dr. R. Reid-Smith at the Public Health Agency of Canada, Guelph, Canada for providing the *Salmonella* Heidelberg isolate; and to the staff members at the Ottawa Laboratory (Fallowfield), Canadian Food Inspection Agency, Ottawa, Canada, including Sandra McIntosh, Sarah Sauvé, Krystal Belanger, and Tessia Berry for their technical assistance in caring and handling animals; Hanhong Dan for his technical assistance in generating DNA standards for the qPCR assays; and Sally Lloy and Danielle Wolfe for their technical assistance in performing the qPCR assays.

## Author Contributions

JG, and ET designed the experiments. JG wrote the manuscript, all co-authors edited and contributed to revisions. GY, and JG carried out the animal experiments and bacterial enumeration and analyzed the bacterial culture data. GY and MC performed PCR for quantitation of resistance genes. CL, SC and MK conducted the 16S rRNA gene amplicon sequencing analysis. EH, PC and MK performed the whole genome sequencing analysis. All authors have read and agreed to the submitted version of the manuscript.

## Data availability statement

All supporting data have been provided within the article or through supplementary data files. One supplementary table and five supplementary figures are available with the online version of this article. The whole genome sequencing data generated for this study can be found in the NCBI BioProject PRJNA1079257.

