## Supplemental figures for "Gut Microbiome Perturbation Affected Conjugative Transfer of Antimicrobial Resistance Genes"

Supplementary table 1. PCR primers and conditions for targeted gene quantification

| Primer | Sequence (5'→3') | Primer (nM) | Annealing (A) & extension (E) | Amplicon size (bp) | Target gene | Reference |
|---|---|---|---|---|---|---|
| BACT1369F<br>PROK1492R | CGGTGAATACGTTCYCGG<br>GGWTACCTTGTTACGACTT | 300 | A: 56 ºC, 30 sec;<br>E: 72 ºC, 30 sec | 123 | small-subunit rRNA gene, *rrnS* | Suzuki *et al.*, 2000 |
| Malo2-F<br>Malo2-Ra | GTATTGTTGATTAATGAGATCCG<br>ATATTACGCACGGAAACACGTT | 250 | A: 55 ºC, 30 sec;<br>E: 72 ºC, 30 sec | 373 | *Salmonella* invasion protein gene, *invA* | Malorny et al., 2001 |
| blaCMY-2F<br>blaCMY-2R | AGG GAA GCC CGT ACA CGT T<br>GCT GGA TTT CAC GCC ATA GG | 300 | A: 52 ºC, 30 sec;<br>E: 72 ºC, 30 sec | 205 | β-lactam resistance gene, $bla_{CMY-2}$ | Boyer & Singer, 2012 |
| blaTEM-1F<br>blaTEM-1R | CATTTTCGTGTCGCCCTTAT<br>GGCGAAAACTCTCAAGGAT | 200 | A: 58 ºC, 30 sec;<br>E: 72 ºC, 30 sec | 167 | β-lactam resistance gene, $bla_{TEM-1}$ | Resende. et al. 2014 |
| intI1-F2<br>intI1-R2 | TCGTGCGTCGCCATCACA<br>GCTTGTTCTACGGCACGTTTGA | 400 | A & E: 62 ºC, 60 sec | 67 | integrase class 1, *intI*1 | Gaze *et al.*, 2011 |
| sul1-F<br>sul1-R | GACTGCAGGCTGGTGGTTAT<br>GAAGAACCGCACAATCTCGT | 200 | A & E: 64 ºC, 60 sec | 105 | sulfonamide resistance gene 1, *sul*1 | Marti *et al.*, 2014 |

| | | | | | | |
|---|---|---|---|---|---|---|
| strA-F | TCAATCCCGACTTCTTACCG | 400 | A & E: 62 ºC, | 126 | aminoglycoside-3″-phosphotransferase gene, *strA* | Walsh *et* |
| strA-R | CACCATGGCAAACAACCATA | | 60 sec | | | *al*., 2011 |
| ermF-F | TCGTTTTACGGGTCAGCACTT | 300 | A & E: 61 ºC, 60 | 182 | erythromycin resistance gene locus F, *ermF* | Knapp *et* |
| ermF-R | CAACCAAAGCTGTGTCGTTT | | sec | | | *al*., 2010 |
| tetQ-F | AGAATCTGCTGTTTGCCAGTG | 500 | A & E: 63 ºC, 60 | 169 | tetQ | Aminov |
| tetQ-R | CGGAGTGTCAATGATATTGCA | | sec | | | *et al.* |
| | | | | | | 2001 |
| cfxA-F | TGACTGGCCCTGAATAATCT | 1000 | A: 55 ºC, 30 sec; | 301 | | Eitel *et* |
| cfxA-R | ACAAAAGATAGCGCAAATCC | | E: 72 ºC, 30 sec | | | *al*., 2013 |

**Supplemental figure legends**

Figure S1. Enumeration of *Salmonella* Typhimurium and *S.* Heidelberg (mean ± SE) in fecal samples from mice following the pre-treatment with no antibiotic (None), ampicillin (Amp), streptomycin (Strep) or sulfamethazine (Sulf); n = 5, 6, 5 and 5 for None, Amp, Strep and Sulf, respectively. ** indicate that the *Salmonella* abundance was significantly ($p$ <0.01) different from those of the other treatment groups within the same serovar on the same day post infection (DPI).

Figure S2. Survival curves of C57BL/6 mice that were inoculated with *Salmonella* Typhimurium and *Salmonella* Heidelberg following the pre-treatment with no antibiotic (blue, n = 5), ampicillin (red, n = 6), streptomycin (green, n = 5), or sulfamethazine (purple, n = 5). Mortality occurred when mice either died unexpectedly or were euthanized due to morbidity. The asterisk (*) indicates $p < 0.05$ when comparing streptomycin (green curve) to other pre-treatments (blue, red or purple curve).

Figure S3. Pearson correlation ($P < 0.0001$) between the quantity of *sul1* (A), *intI1* (B), and *strA* (C) genes and the number of *Salmonella* in individual mouse fecal samples from all mice in this study with best fit line (solid) and 95% confidence bands (dotted).

Figure S4. Microbial species richness as assessed by the Chao1 index on each day post infection (DPI) from various per-treatment groups: None = no antibiotic, Amp = ampicillin, Strep = streptomycin, Sulf = sulfamethazine.

Figure S5. Principal coordinate analysis (PCoA) on Bray-Curtis dissimilarity of mouse gut microbiota derived from individual mice on each day post infection (DPI) from various pre-treatment groups: None = no antibiotic, Amp = ampicillin, Strep = streptomycin, Sulf = sulfamethazine.

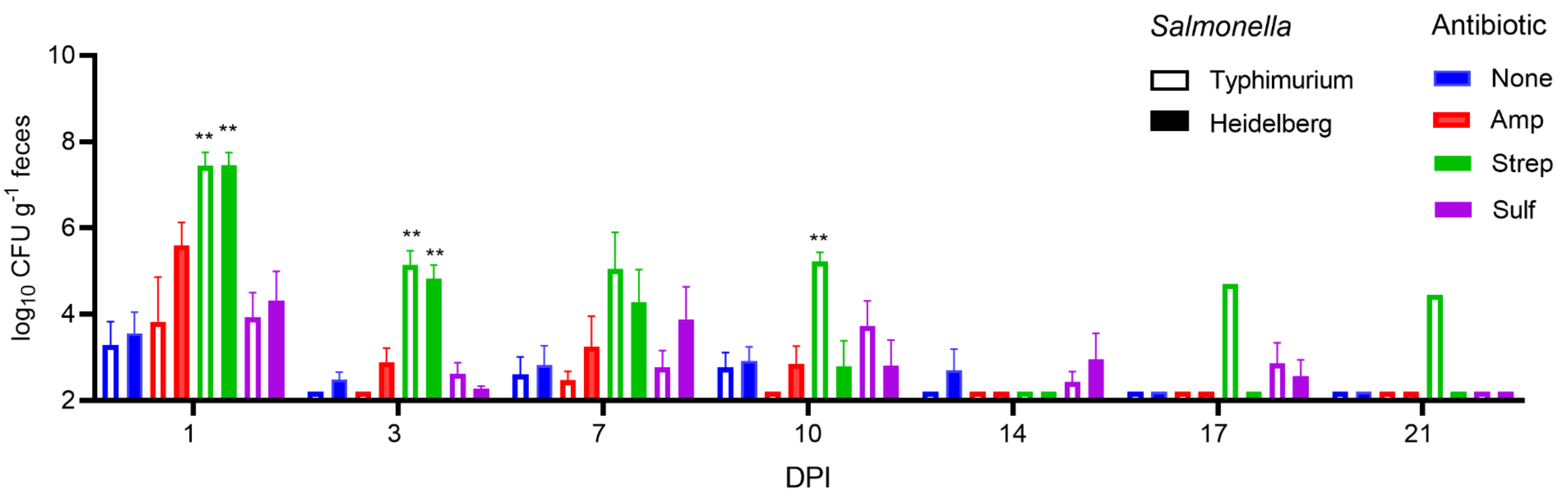
Salmonella
Typhimurium
Heidelberg
Antibiotic
None
Amp
Strep
Sulf
log10 CFU g-1 feces
10
8
6
4
2
**
1
3
7
10
14
17
21
DPI

Figure S1

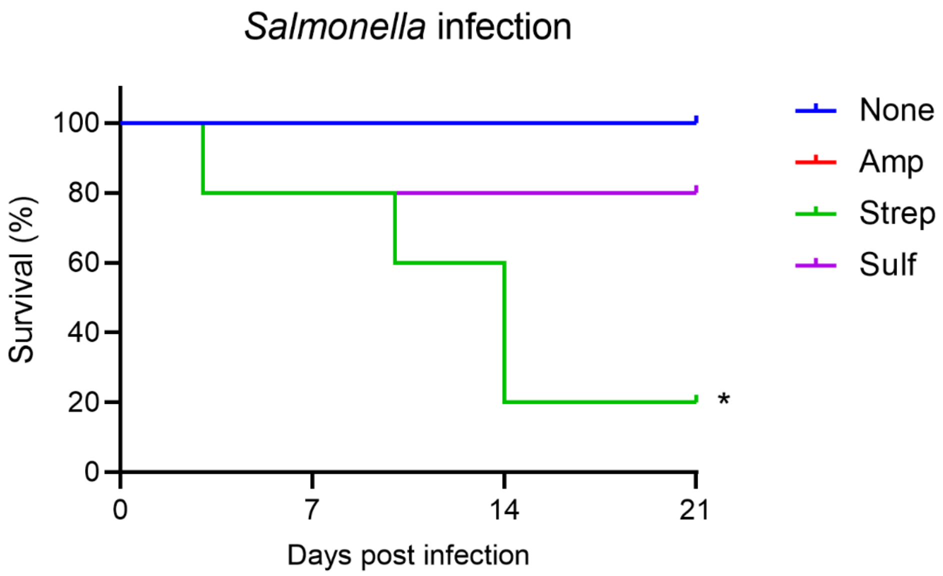

Salmonella infection
Survival (%)
100
80
60
40
20
0
0
7
14
21
Days post infection
*
None
Amp
Strep
Sulf


Figure S2

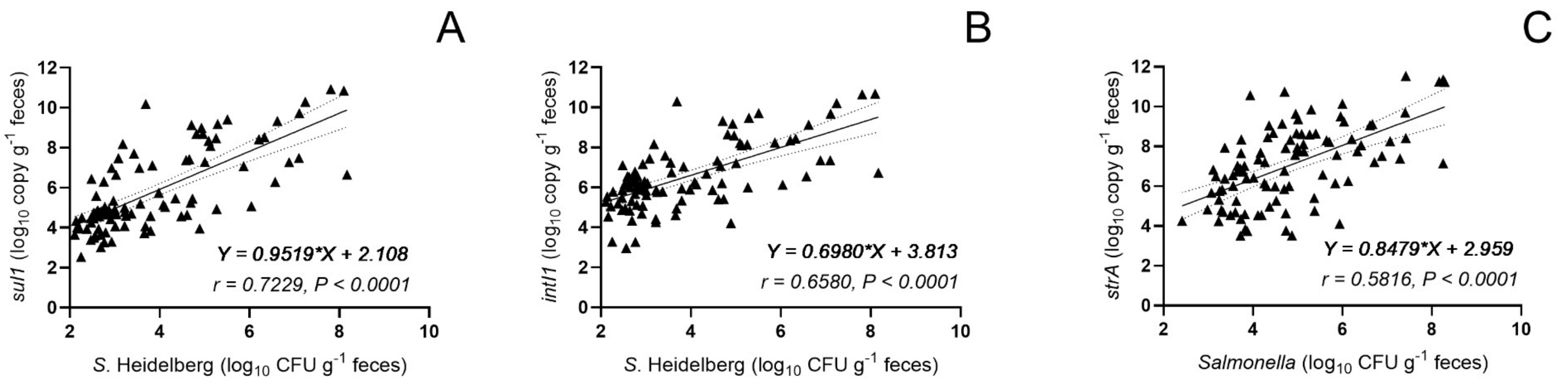
A
B
C
sul1 (log10 copy g-1 feces)
intI1 (log10 copy g-1 feces)
strA (log10 copy g-1 feces)
S. Heidelberg (log10 CFU g-1 feces)
Salmonella (log10 CFU g-1 feces)
Y = 0.9519*X + 2.108
r = 0.7229, P < 0.0001
Y = 0.6980*X + 3.813
r = 0.6580, P < 0.0001
Y = 0.8479*X + 2.959
r = 0.5816, P < 0.0001

Figure S3

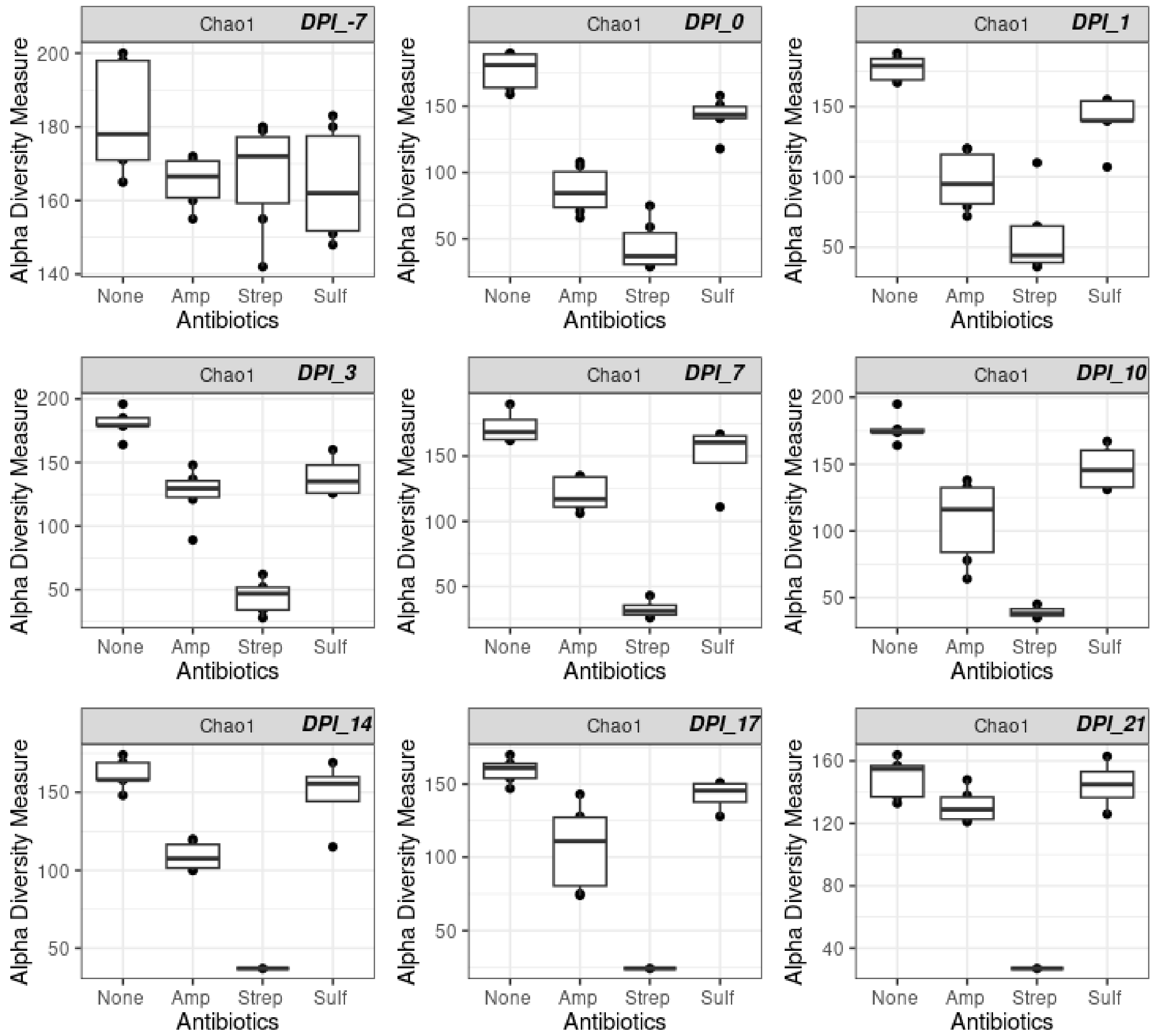
Chao1
DPI_-7
DPI_0
DPI_1
DPI_3
DPI_7
DPI_10
DPI_14
DPI_17
DPI_21
Alpha Diversity Measure
Antibiotics
None
Amp
Strep
Sulf

Figure S4

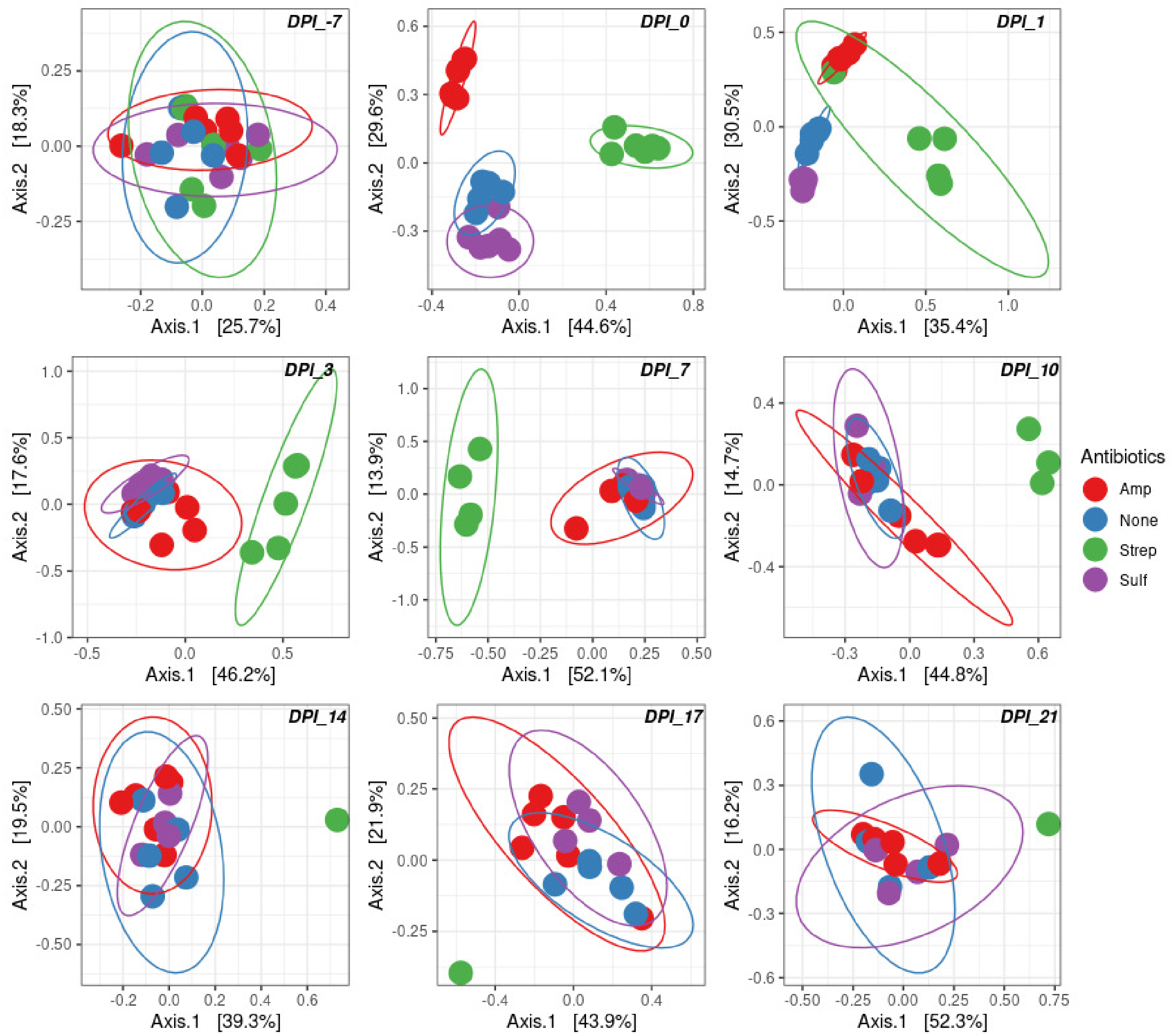
DPI_-7
DPI_0
DPI_1
DPI_3
DPI_7
DPI_10
DPI_14
DPI_17
DPI_21
Axis.1 [25.7%]
Axis.2 [18.3%]
Axis.1 [44.6%]
Axis.2 [29.6%]
Axis.1 [35.4%]
Axis.2 [30.5%]
Axis.1 [46.2%]
Axis.2 [17.6%]
Axis.1 [52.1%]
Axis.2 [13.9%]
Axis.1 [44.8%]
Axis.2 [14.7%]
Axis.1 [39.3%]
Axis.2 [19.5%]
Axis.1 [43.9%]
Axis.2 [21.9%]
Axis.1 [52.3%]
Axis.2 [16.2%]
Antibiotics
Amp
None
Strep
Sulf

Figure S5
